# Harnessing Ionic Selectivity In Acetyltransferase Chemoproteomic Probes

**DOI:** 10.1101/2020.09.24.311530

**Authors:** Yihang Jing, Jose Montano, Michaella Levy, Jeff Lopez, Pei-Pei Kung, Paul Richardson, Krzysztof Krajewski, Laurence Florens, Michael Washburn, Jordan L. Meier

## Abstract

Chemical proteomics provides a powerful strategy for the high-throughput assignment of enzyme function or inhibitor selectivity. However, identifying optimized probes for an enzyme family member of interest and differentiating signal from background remain persistent challenges in the field. To address this obstacle, here we report a physiochemical discernment strategy for optimizing chemical proteomics based on the Coenzyme A (CoA) cofactor. First, we synthesize a pair of CoA-based Sepharose pulldown resins differentiated by a single negatively charged residue, and find this change alters their capture properties in gel-based profiling experiments. Next, we integrate these probes with quantitative proteomics and benchmark analysis of ‘probe selectivity’ versus traditional ‘competitive chemical proteomics’. This reveals the former is well-suited for the identification of optimized pulldown probes for specific enzyme family members, while the latter may have advantages in discovery applications. Finally, we apply our anionic CoA pulldown probe to evaluate the selectivity of a recently reported small molecule N-terminal acetyltransferase inhibitor. These studies further validate the use of physical discriminant strategies in chemoproteomic hit identification and demonstrate how CoA-based chemoproteomic probes can be used to evaluate the selectivity of small molecule protein acetyltransferase inhibitors, an emerging class of pre-clinical therapeutic agents.

## Introduction

Over the past two decades, chemical proteomics (also referred to as ‘chemoproteomics’) has emerged as a powerful strategy for the high-throughput annotation of protein function.^*1*^ This approach involves using an active site probe to capture an enzyme class of interest from cells or a proteomic sample, allowing the biological activity of multiple enzymes within that family to be assessed in parallel.^*2*^ The ability of synthetic small molecules or metabolites to compete chemoproteomic capture can provide a rapid readout of drug target engagement or ligand selectivity across several related enzyme family members.^*3*^ Chemoproteomic profiling strategies have been applied to a wide-range of enzyme families including hydrolases,^*4, 5*^ metalloenzymes,^*6*^ kinases,^*7, 8*^ methyltransferases,^*9*^ and deacylase enzymes.^*9–12*^ Recently, in the process of characterizing a histone acetyltransferase chemoproteomic probe, we discovered that CoA-based affinity resins exhibit broad-spectrum capture properties, enriching lysine acetyltransferases, N-terminal and metabolic acetyltransferases, and several CoA and NAD-based metabolic enzymes (e.g. oxidoreductases).^*13, 14*^ This indicates CoA-based chemoproteomic probes may have utility in studying the pharmacological interactions of additional enzyme classes, beyond histone acetyltransferases. However, one challenge in applying this approach is how to differentiate a ‘true hit’ from background. Many histone acetyltransferases utilize an ordered bi- bi- binding mechanism, with acetyl-CoA binding first, which allows their chemoproteomic capture to be competed by pre-incubating lysates with acetyl-CoA.^*15*^ However, for acetyltransferases that don’t exhibit this binding mode, or which bind to a different ligand (e.g. crotonyl-CoA, NAD), acetyl-CoA competition may not be evident, limiting our ability to discern selective chemoproteomic enrichment from noise.

An alternative approach to distinguishing signal from noise in chemoproteomic experiments is to compare how enzymes are enriched by two physiochemically distinct capture probes. This was compellingly demonstrated in a recent study, where a comparison of proteins captured by clickable photoaffinity probes differing only in their absolute stereochemistry at a single molecular recognition determinant streamlined the identification of selective protein-fragment interactions.^16^ Inspired by this approach, we hypothesized that a similar physiochemical discernment strategy could be employed to identify new targets selectively enriched by CoA-based probes, thus expanding the scope of protein-ligand interactions open to chemoproteomic interrogation.

## Results

To test this hypothesis, we designed a pair of CoA-based chemoproteomic capture resins distinguished by only a single amino acid replacement (Fig. 1a). Capture resin **1** consists of CoA tethered through a thioether linkage to a series of four 6-aminohexanoic acid (Ahx) linkers. Similar to Lys-CoA, Ahx-CoA analogues have been found to bind a wide variety of acetyltransferases with good affinity while also providing a straightforward handle for functionalization, enabling them to be used as a general scaffold for the development for fluorescence polarization assays.^*16, 17*^ In addition to facilitating our physiochemical discernment strategy (vide infra), we hypothesized characterizing their proteome-wide binding properties might expand the applications of these novel CoA analogues. Capture resin **2** is near identical in structure, containing a single substitution of the first Ahx residue (adjacent to the CoA) with an aspartate. Our rationale was since many acetyltransferases bind to positively charged substrates (such as histone tails and protein N-termini), incorporation of a negative charge near the active site may serve as an effective molecular recognition discriminant or, alternatively, facilitate the selective enrichment of acetyltransferases that recognize negatively charged substrates such as amino acids and RNA.^*18, 19*^ Based on this design, we synthesized two CoA analogues, CoA-(Ahx)4 or CoA-D-(Ahx)3, using a previously reported route^*20*^ from CoA and the cognate bromoacetamide peptide. Following HPLC purification and quantification, the two CoA analogues were separately coupled to NHS-Sepharose via the epsilon amine of a C-terminal lysine residue, giving access to affinity resins **1** and **2**.

**Figure 1.**
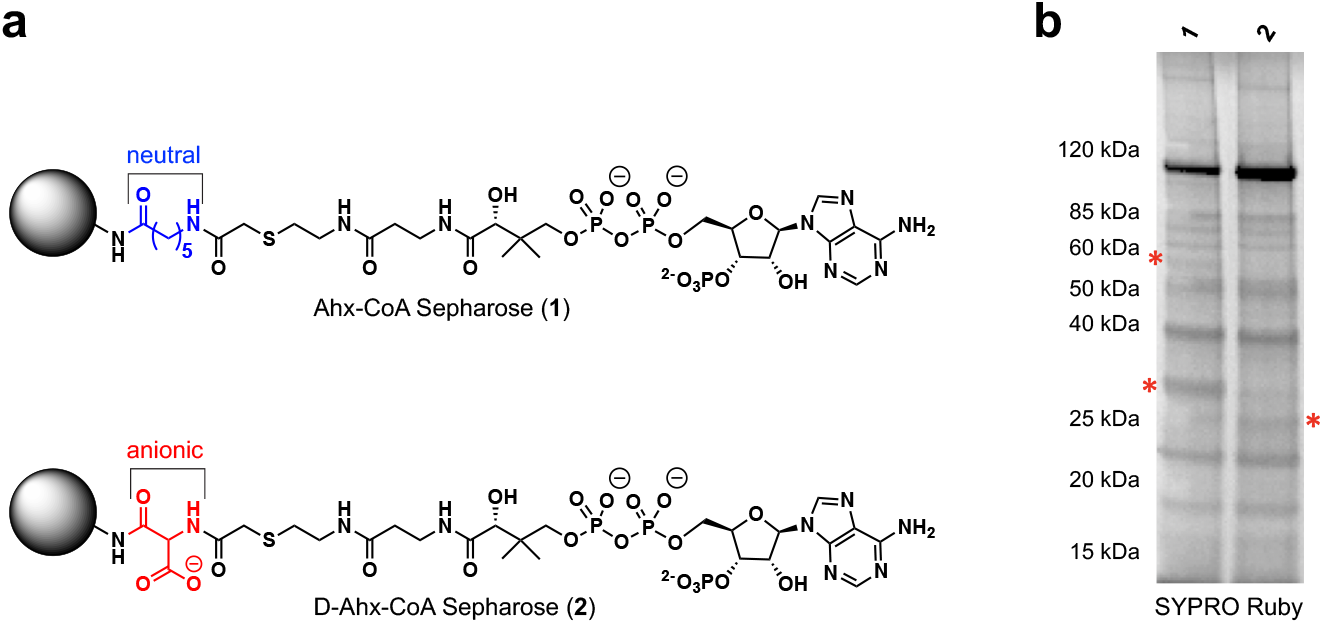
Exploring ionic selectivity in acetyltransferase chemoproteomic probes. (a) Structure of general acetyltransferase capture resins applied in this study, emphasizing the differing neutral (blue) and anionic (red) residues introduced as physical discriminants. Both molecules are tethered to the resin by Ahx-Ahx-Ahx-Lys linkers (structure not shown; see Supporting Information for synthetic details). (b) Gel-based analysis of proteins enriched by chemoproteomic resins **1** and **2**, as assessed by SYPRO Ruby staining of captured and eluted proteins. Arrows indicate positions of differentially enriched targets.

As an initial test of the selectivity of **1** and **2**, we quantitatively examined their overall proteomic interaction profiles using SDS-polyacrylamide electrophoresis (SDS-PAGE)-based profiling. Briefly, freshly prepared proteomes from chronic myelogenous leukemia (K562) cells were incubated with resins **1** and **2** at room temperature for one hour. Resins were then washed, eluted, subjected to SDS-PAGE, and incubated with SYPRO Ruby to facilitate visualization by in gel fluorescence scanning. The intensity of the proteomic enrichment profiles by resins **1** and **2** are overall similar, consistent with CoA-Sepharose binding being a major determinant of capture (Fig. 1b). Many of the observed interactions were strongly competed with acetyl-CoA (0.1 mM), as has been observed in previous studies (Fig. S1). However, encouragingly, a subset of proteins showed signs of selective enrichment by charged and uncharged resins (Fig. 1b, red asterisks). This suggests a protein subset may be preferentially captured by the physiochemically distinct capture resins, prompting us to determine their identities using quantitative LC-MS/MS proteomics.

To facilitate the quantitative detection of proteins binding to **1** and **2**, we integrated our previously reported chemoproteomic enrichment and mass spectrometry–based proteomic protocol with a post-enrichment derivatization step using Tandem Mass Tags (TMT) (Fig. 2a).^*21*^ In this approach, samples are enriched by **1** or **2**, subjected to tryptic digest, labeled in parallel with a set of isobaric mass tags with a single isotopic substitution, and pooled together. Tagged peptides from different samples exhibit indistinguishable migration and mass to charge ratios during MS1 analysis, but yield distinct mass reporter ions upon MS2 fragmentation. The relative intensity of these mass reporter ions provides a measure of relative peptide (and parent protein) abundance found in each enriched sample. Of note, the use of TMT tagging represents a technical advance over our previous studies, which have demonstrated the utility of label-free spectral counting in analysis of acyl-CoA/protein interactions.^*14*^ Applying our chemoproteomic probes in combination with a 4-plex TMT-tagging strategy allowed comparative quantification of proteins enriched from K562 proteomes under four affinity capture conditions: i) probe **1** (no competitor), ii) probe **2** (no competitor), iii) probe **1** (0.1 mM acetyl-CoA competitor), and iv) probe **2** (0.1 mM acetyl-CoA competitor; Fig. 2a). Three independent experiments were performed, and individual protein abundances were normalized relative to the overall protein intensity quantified in each tagged sample in order to account for any technical variations (i.e. sample loss) introduced prior to TMT-tagging. For these studies, a ‘selective interaction’ with a chemoproteomic affinity resin was operationally defined as proteins found to be preferentially enriched by an average value of >2 fold by **1** or **2**, which is similar to the criteria used in previous studies.^*16*^

**Figure 2.**
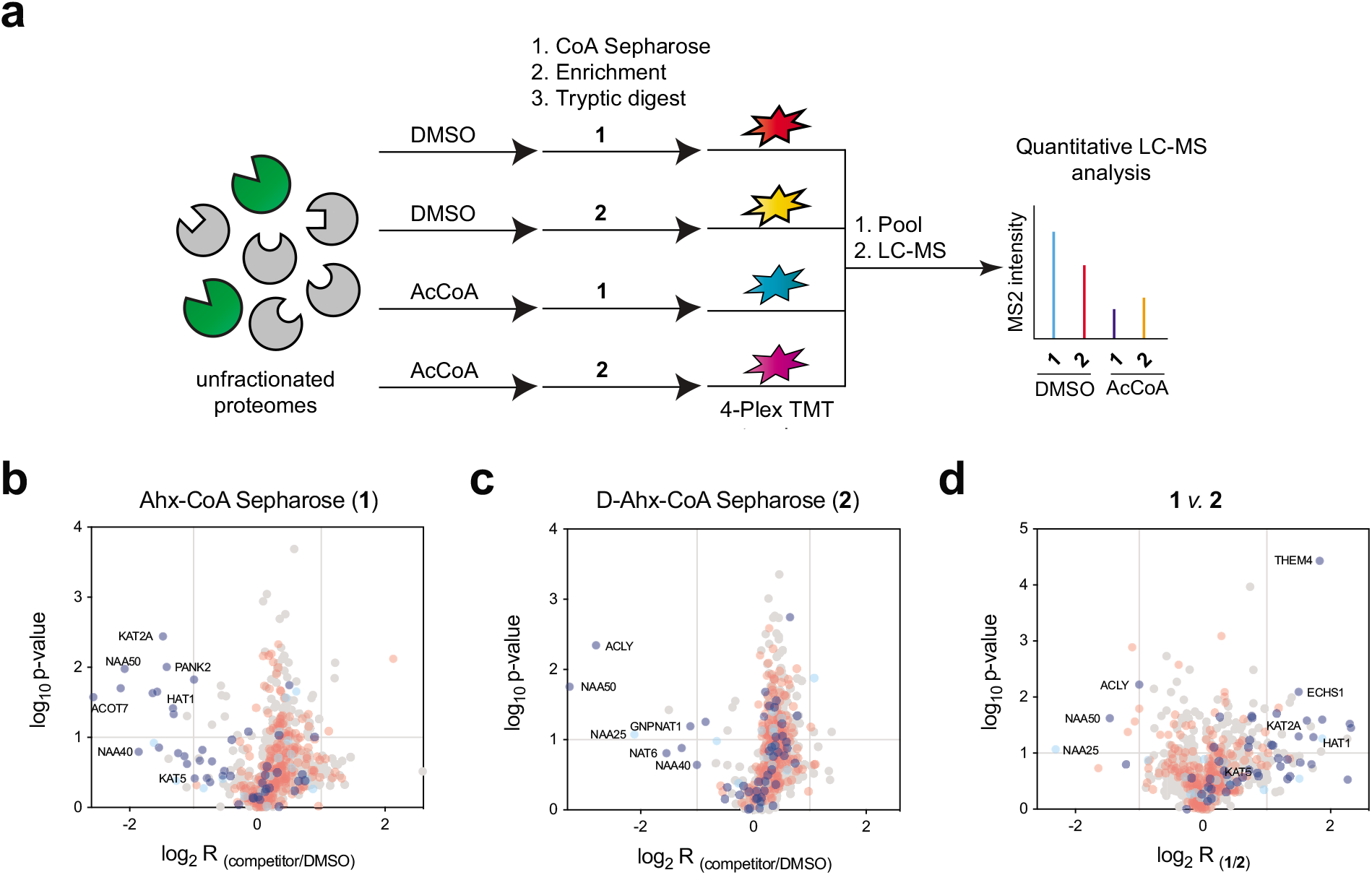
Proteomic targets of neutral and anionic acetyltransferase chemoproteomic probes. (a) General workflow for chemoproteomic capture and Tandem Mass Tag (TMT)-based LC-MS/MS quantification of enriched proteins. (b) Volcano plot of proteins enriched by resin **1** versus acetyl-CoA (0.1 mM) competition. (c) Volcano plot of proteins enriched by resin **2** versus acetyl-CoA (0.1 mM) competition. (d) Volcano plot of proteins enriched by resin **1** versus resin **2**. Dark blue = known CoA-binding proteins, light blue = known acetyltransferase interactors, red = known Sepharose resin-binding proteins, gray = other.

Applying this methodology, we identified 71 proteins that displayed selective interaction with probe **1**, and an additional 17 that selectively interacted with anionic probe **2** (Table S1). For comparison, we exploited TMT multiplexing to simultaneously analyze a competitive chemoproteomic experiment, examining the ability of acetyl-CoA (0.1 mM) to block enrichment of proteins by **1** and **2**. Here we identified 32 and 15 proteins, respectively, whose capture by **1** and **2** was competed >2-fold by acetyl-CoA. To better understand what types of proteins selectively interact with resins **1** and **2**, we used literature data to classify identified targets into three categories: direct CoA binders (e.g. Uniprot-annotated acetyltransferases; dark blue), indirect interactors (e.g. members of acetyltransferase complexes; light blue), or non-specific Sepharose-binding proteins (compiled from a recent proteomic dataset;^*14*^ red). Overlaying these annotations onto our competitive chemoproteomic dataset revealed a clear proteomic subset enriched in CoA-binding proteins and acetyltransferase interactors (that was efficiently enriched by **1** and **2** and potently competed by acetyl-CoA (Fig. 2b and Fig. 2c, top left). In contrast, proteins known to be non-specifically enriched by the Sepharose matrix (red) were mostly found clustered in the center of graph, indicating their capture was not competed by acetyl-CoA. To assess whether our ionic discriminant was able to identify CoA-binding proteins and AT interactors with similar efficacy, we graphed the relative enrichment of proteins by **1** versus **2** (Fig. 2d). In this analysis two disparate clusters of proteins enriched in CoA binders and AT interactors can be visualized, one which shows preferential enrichment by **1** (Fig. 2d, top right) and one which shows preferential enrichment by **2** (Fig. 2d, top left). This suggests that employing physiochemical discriminants may offer a valuable complement to competitive chemoproteomics by distinguishing authentic hits (in this case CoA binders and AT interactors) from background while simultaneously specifying optimized probes to interrogate their pharmacological function.

To validate the selective enrichment of acetyltransferases by **1** and **2**, we performed pulldowns and monitored protein levels by western blotting. Consistent with our chemoproteomic findings, we found neutral probe **1** provided greater enrichment of KAT2A, HAT1, and KAT8, while anionic probe **2** demonstrated more efficient enrichment of NAA10, NAA25, NAA80, ACLY, and NAA50 (Fig. 3a, S2). Enrichment of these acetyltransferases was also efficiently competed by preincubation of lysates with acetyl-CoA, further confirming their specific capture. To understand the physiochemical basis for the selective enrichment of specific protein acetyltransferases by **1** and **2**, we surveyed their known biochemical substrates reported in the literature (Fig. 3b). For example, capture resin **1** selectively enriches KAT2A and HAT1 (>2-fold), both of which acetylate positively charged histone tails. This selective capture may reflect the fact that bisubstrate inhibitor **2** places a negative charge in the substrate binding site, a change which may be reasonably expected to reduce the affinity for these enzymes. In contrast, protein acetyltransferases found to be selectively enriched (>2-fold) by capture resin **2** include NAA10 and NAA80. Each of these enzymes recognize and modify peptides with acidic N-termini (Fig. 3b), and their enhanced capture by probe **2** may reflect interaction of its aspartate residue with positively charged active site residues in a manner that mimics this interaction (Fig. 3c). Besides protein acetyltransferases, probe **2** selectively enriched additional CoA-binding proteins that interact negatively charged substrates, including ACLY which binds to CoA and citrate, and GNPNAT, which acetylates the anionic metabolite glucosamine-6-phosphate. An exception to this trend is the preferential capture of ATAT1 by neutral probe **1**, as ATAT1 recognizes a tubulin lysine substrate with a flanking aspartate residue. This latter observation highlights the complementary importance of design and empirical determination of relative capture efficiencies to design optimized affinity resins. Overall, these studies suggest partial substrate mimicry as a biochemical rationale for the ability of ionic charge to act as a physiochemical discriminant in CoA-based chemoproteomic probes.

**Figure 3.**
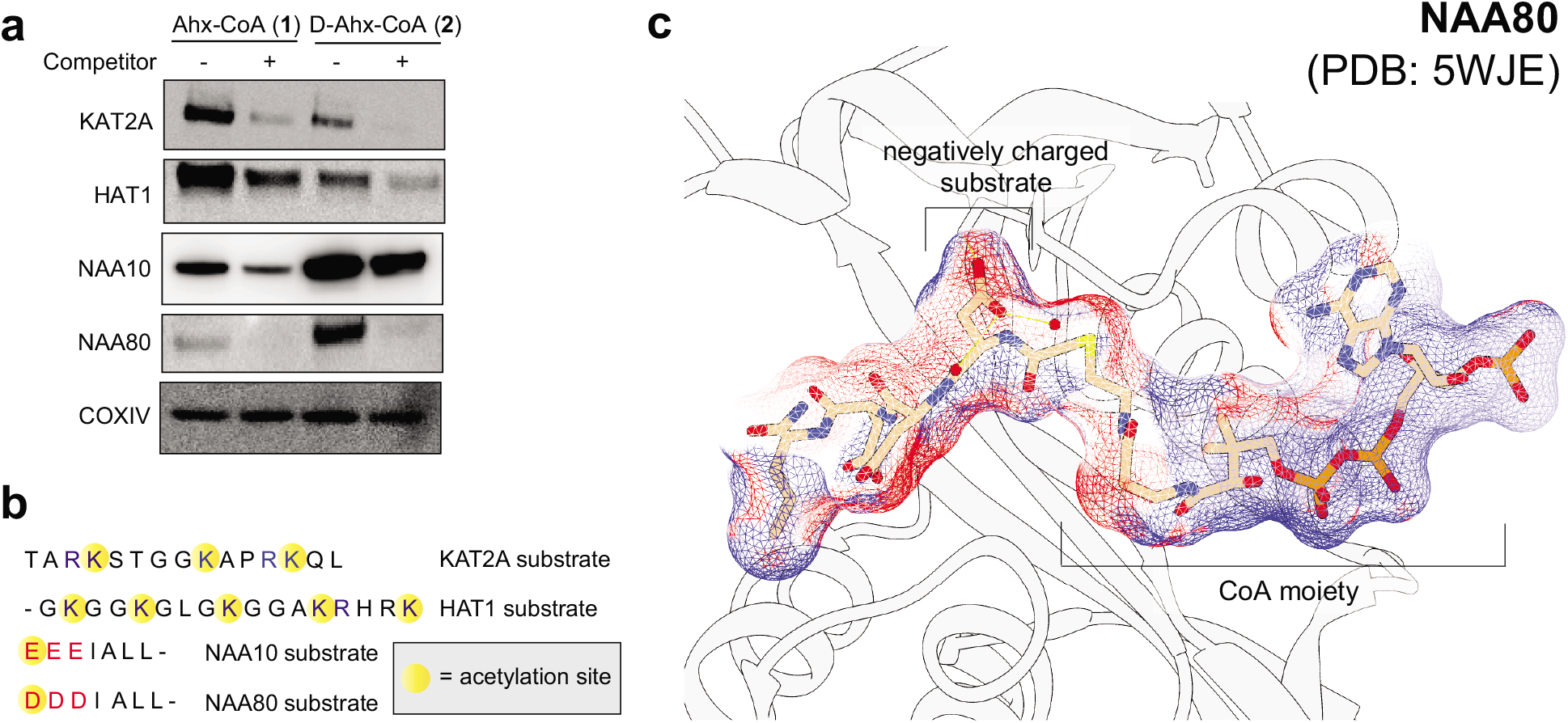
Assessing the differential enrichment of acetyltransferases by ionic and non-ionic probes. (a) Validation of differential enrichment of KAT2A, HAT1, NAA10, and NAA80 by resins **1** and **2**. (b) Amino acid sequences of known biochemical substrates of lysine and N-terminal protein acetyltransferases known in literature. Dashes equal truncated sequences. (c) Crystal structure showing accommodation of negatively charged bisubstrate-CoA ligand in acetyltransferase active site of a NAA80 homologue (PDB: 5WJE). Density map surrounding the bisubstrate ligand represents electrostatic potential calculated using the APBS-PDB2PQR software suite and visualized in Chimera.

Finally, we sought to apply our knowledge of chemoproteomic probe design to better understand the pharmacology of acetyltransferase inhibitors. Recently, we reported a synthetic small molecule probe of the N-terminal acetyltransferase NAA50, an essential human gene that is critical for sister chromatid cohesion and chromosome condensation.^*22*^ This small molecule (referred to here as compound **3**; Fig. 4a) was derived from medicinal chemistry optimization of a hit from a DNA-encoded library screen, and binds NAA50 with low nanomolar affinity in biochemical and competitive chemoproteomic analyses. However, one caveat to the chemoproteomic assays used in this initial characterization was the use of a Lys-CoA-based resin which provides sub-optimal enrichment of N-terminal acetyltransferases. This impeded analysis of the proteomic selectivity with which **3** binds to NAA50 compared other to other members of the N-terminal acetyltransferase family. To address this shortcoming we applied anionic probe **2**, which preferentially captures N-terminal acetyltransferases due to mimicry of their negatively charged substrates, to analyze the selectivity of **3** in proteomic extracts using immunoaffinity profiling.^*13*^ In this experiment, cell lysates form K562 cells were incubated with increasing amounts of inhibitor **3** or vehicle DMSO, subjected to chemoproteomic capture using probe **2**, and probed with an anti-N-terminal acetyltransferase antibody (Fig. 4b). Reduced antibody signal in the presence of ligand indicates blockade of enrichment, and is a sign of a ligand-acetyltransferase interaction. Using this approach we found that pre-incubation of proteomes with small molecule inhibitor **3** followed by chemoproteomic capture led to a dose-dependent reduction in NAA50 capture, consistent with our prior chemoproteomic studies (Fig. 4c). Applying the same approach to probe NAA10, which represents one of the most predominant and well-studied cellular N-terminal acetyltransferases,^*23*^ we observed minimal competition. Interestingly, NAA10 and NAA50 are known to co-occupy the NatA complex.^*13, 24*^ The lack of competition of NAA10 in our experiment suggests our chemoproteomic resin **2** does not indirectly enrich NAA10 through the NAA50-containing NatA complex, but rather reports on the active-sites of these two different N-terminal acetyltransferases independently. To examine the interaction of inhibitor **3** with an N-terminal acetyltransferase that does not co-occupy a protein complex with NAA50, we turned our attention to NAA20 and NAA80, two of the proteins most differentially-enriched by anionic capture resin **2**.^*25–27*^ No antibody for NAA20 available commercially, which led us to monitor enrichment of NAA25 (a member of NAA2O-containing NatB complex) as a proxy. In this case we also observed essentially no competition of capture, again indicating the high selectivity of **3** for NAA50. These studies demonstrate how the optimization of chemoproteomic profiling probes using a physical discriminant approach can facilitate analysis of synthetic acetyltransferase inhibitors in complex proteomic contexts.

**Figure 4.**
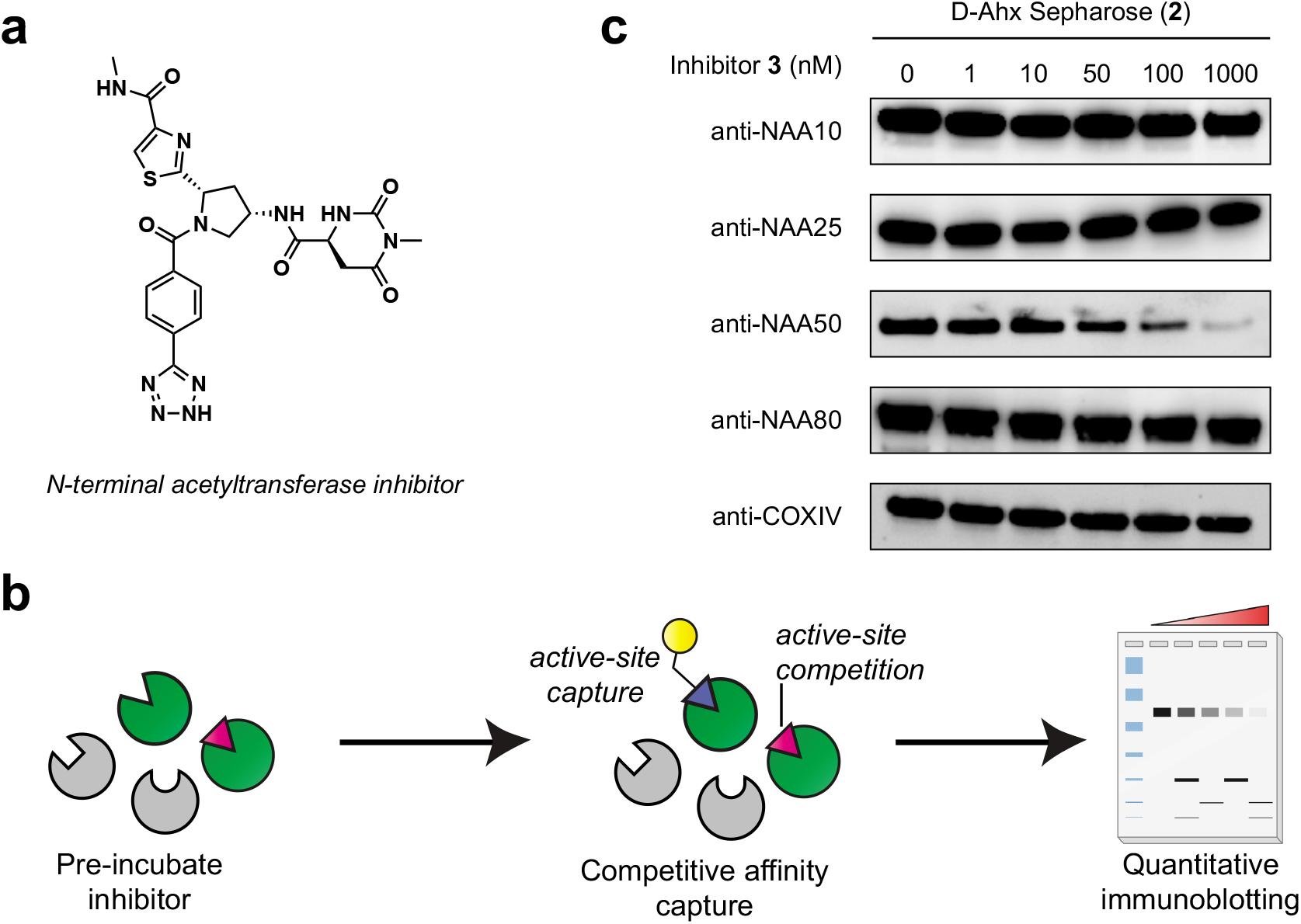
Applying anionic chemoproteomic capture resins to study the selectivity of a synthetic N-terminal acetyltransferase inhibitor. (a) Structure of Naa50 inhibitor. (b) Scheme for competitive immunoaffinity profiling experiment. Western blot illustration by Biorender. (c) Competitive chemoproteomic analysis of inhibitor’s interaction with Naa10, Naa20 (Naa25), Naa50, and Naa80.

## Discussion

Here we have reported the utilization of ionic selectivity as a physical discriminant in the capture of acetyltransferases by chemoproteomic probes. We find that incorporation of a single negative charge can facilitate the identification of known CoA binders and AT interactors, in particular differentiating epsilon lysine and N-terminal protein acetyltransferases. This provides a useful complement to competitive chemoproteomic profiling methods by simultaneously enriching specific molecular interactions from background and specifying optimized probes for pharmacological profiling. An unexpected finding of this study was that several acetyltransferases are preferentially enriched by anionic probe **2**, which was designed to function control to differentiate histone acetyltransferases from background. In the future, further elaboration of anionic CoA-based probes may enable more sensitive or specific proteomic interrogation of the N-terminal acetyltransferase enzyme class. Negatively charged substrate-based CoA inhibitors of these proteins have been previously reported, and represent a promising lead for the design of such agents.^*28*^ In addition to acetyltransferases, we identified several CoA binders and AT interactors that show selective enrichment by probes **1** or **2**. While we caution that the direct interaction of our probes with many of these targets remains to be validated, a recent study has demonstrated that similar Ahx-CoA analogues can be readily converted into fluorescence polarization probes for small molecule inhibitor screening.^*17*^ This suggests the potential for the chemoproteomic probe-enzyme pairs described to may provide a universal entry into biological profiling, functional enzyme analysis, and inhibitor development, as has been demonstrated for other probe classes.^*29–31*^ Finally, we note several limitations of our approach as currently constituted. In contrast to previous studies of clickable photoaffinity probes that used enantiomeric probe pairs as a physical discriminant,^*16*^ our CoA-based probes are unable to clearly distinguish direct and indirect interactors due to the non-covalent nature of pulldown method. In the future, this may be addressed by integrating photocrosslinking and stringent washing into this approach.^*32*^ In addition, we found that our overall coverage of AT interactors and CoA-binding proteins in these experiments was shallower relative to some of our recent studies, potentially reflecting a loss of ions due to the relatively long MS2 cycling times required for TMT-based quantification. We anticipate future applications of this technique may benefit from the use of increased chromatographic separation or next-generation LC-MS/MS mass spectrometry platforms. While we have demonstrated the ability of our physical discriminant approach to identify chemoproteomic probes that capture acetyltransferases with higher affinity, one last consideration is that for competitive chemoproteomic experiments such as our analysis of the NAA50 inhibitor to be effective, the affinity of the capture probe and competitor must be roughly matched. In other words, if a chemoproteomic capture agent binds too tightly to a protein of interest, no competition will be observed. While the absolute affinity of **1** and **2** for the acetyltransferases enriched in this study have not been rigorously determined, other groups have reported that dissociation constants of chemoproteomic affinity resins for protein targets can be estimated by incorporating measurements of target depletion from whole cell lysates.^*33*^ Similar approaches may be important in applications of CoA-based probes to study the proteome-wide selectivity of small molecule ligands. Drug-like protein acetyltransferases inhibitors have recently been developed and shown promise in oncology and other clinical settings^*34–37*^ Methods to assess target occupancy and ligand selectivity such as those demonstrated here will likely be critical to the continued development and therapeutic validation these pre-clinical agents.

## Supporting Information

Supporting Information including Figures S1–2, Supplementary Table S1, supporting materials, and methods is available online. Mass spectrometry proteomics data has been deposited to the ProteomeXchangeConsortium via the PRIDE partner repository with the dataset identifier PXD020856 and 10.6019/PXD020856

Reviewer account information:

**Password:** BdHP7ngW

## Acknowledgments

This work was supported by the Intramural Research Program of the National Institutes of Health, the National Cancer Institute, The Center for Cancer Research (ZIA BC011488-06), the Stowers Institute for Medical Research, and the National Institute of General Medical Sciences of the National Institutes of Health (RO1GM112639 to MPW).

