## Supplemental Information for "Harnessing Ionic Selectivity In Acetyltransferase Chemoproteomic Probes"

*Yihang Jing,^[a]#^ Jose Montano,^[a]#^ Michaela Levy,^[b]^ Jeffrey Lopez,^[a]^ Pei-Pei Kung,^[c]^ Paul Richardson,^[c]^ Kryzjtof Krajewswki,^[d]^ Laurence Florens,^[b]^ Michael Washburn,^[b],[e]^ and*

*Jordan L. Meier*^[a]^*

^[a]^ Chemical Biology Laboratory, Center for Cancer Research, National Cancer Institute, Frederick, MD, USA. ^[b]^ Stowers Institute for Medical Research, Kansas City, MO, USA. ^[c]^ Worldwide Research and Development, Pfizer Inc., San Diego, CA, USA. ^[d]^ Department of Biochemistry and Biophysics, The University of North Carolina, Chapel Hill, NC, USA. ^[e]^ Department of Pathology and Laboratory Medicine, University of Kansas Medical Center, Kansas City, KS, USA.

**Table of Contents for Supporting Information**

**Page**

Table of Contents	S1

Supplementary Figures S1-S2	S2-3

General materials and methods	S4

Preparation of CoA Sepharose resins	S5

TMT labeling LC-MS/MS analysis	S6

Chemical capture and immunoaffinity profiling protocol	S7

Full gels and blots	S8-18

References	S19

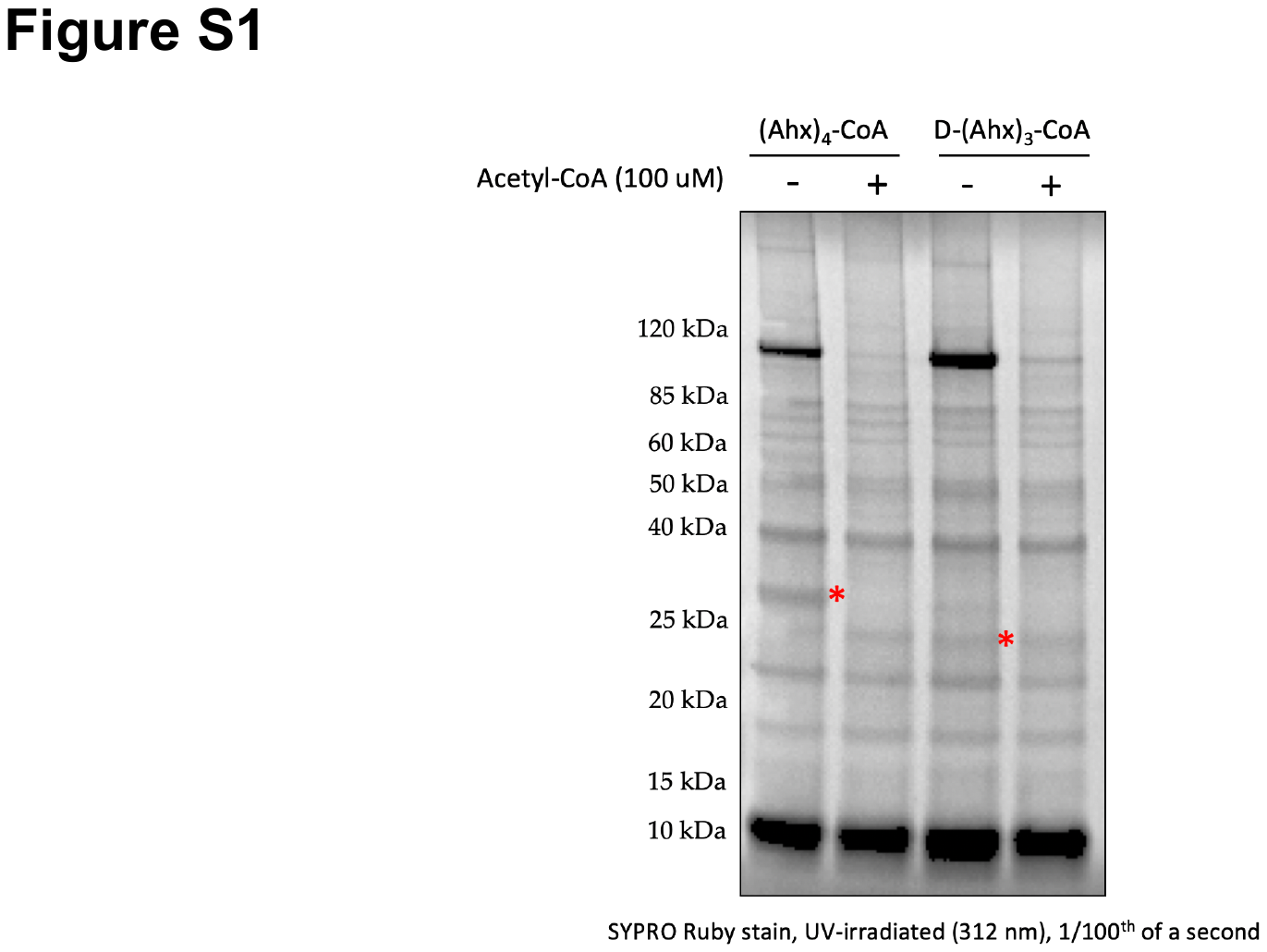

**Figure S1.** Competitive profiling data for enrichment of proteins by chemoproteomic resins **1** and **2** in the presence of 100 μM acetyl-CoA. Arrows indicate positions of differentially enriched targets.

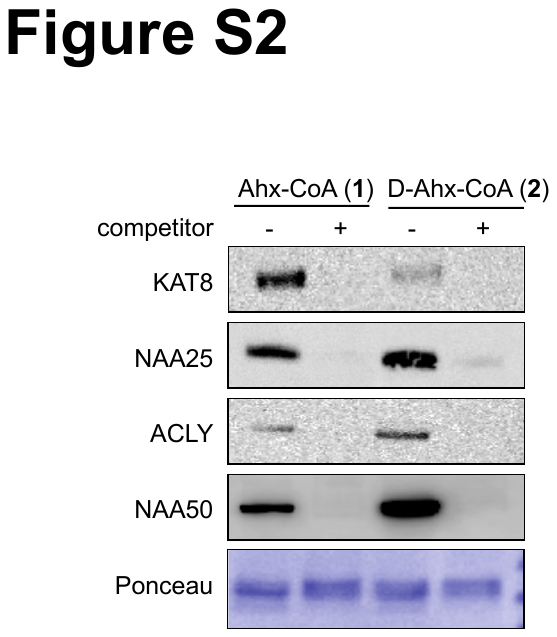

**Figure S2.** Western blot data suggests the selective enrichment of KAT8, NAA25, ACLY, and NAA50 by chemoproteomic resins **1** and **2.**

**General materials and methods**

All solvents used are dried and distilled by using standard methods, and all the chemical reagents and amino acids for peptide synthesis without further noted were purchased from Sigma-Aldrich and Novabiochem. HEK-293T cells were obtained from the NCI Tumor Cell Repository. Primary antibodies: anti-COXIV (#4850), anti-KAT2A (#3305), anti-NAA10 (#13357), and anti-ACLY (#4332) antibodies were purchased from Cell Signaling Technology; anti-NAA25 (ab79490) and anti-NAA80 (ab96844) antibodies were purchased from Abcam; anti-HAT1 (GTX110643) antibody was purchased from GeneTex; anti-KAT8 (A300-992A-T) antibody was purchased from Bethyl Laboratories; anti-NAA50 (16120-1-AP) antibody was purchased from Protein Tech. Secondary antibody: Anti-rabbit IgG, HRP-linked Antibody (#7074) was purchased from Cell Signaling Technology. The peptide used in this research was synthesized on Nova PEG Rink Amide resin (Novabiochem, 855047) following standard Fmoc-based solid-phase peptide synthesis protocol. After the coupling of all amino acids, the removal of protecting groups and cleavage of peptides from the resin were done by incubating the resin with cleavage cocktail containing 95% trifluoroacetic acid (TFA), 2.5% triisopropylsilane, and 2.5% water for 3 hrs. Peptides were purified on a preparative high performance liquid chromatography (HPLC) system, and the mobile phases are buffer A (100% water, 0.1% TFA) and buffer B (90% ACN in water, 0.1% TFA).The purity (> 95%) and identity of peptide was confirmed by analytical LC-MS. For western blotting, SDS-PAGE was performed using Bis-Tris NuPAGE gels (4-12%, Invitrogen #NP0322), and MES running buffer (Life technologies #NP0002) in Xcell SureLock MiniCells (Invitrogen) according to the manufacturer’s instructions. Gels were transferred to nitrocellulose membranes (Novex, Life Technologies #LC2001) by electroblotting at 30 volts for 1 hr using a XCell II Blot Module (Novex). Membranes were blocked using StartingBlock (PBS) Blocking Buffer (Thermo Scientific) for 30 minutes, then incubated overnight at 4°C in a solution containing the primary antibody of interest in the above blocking buffer with 0.05% Tween 20. The membranes were next washed with TBST buffer, and incubated with a secondary HRP-conjugated antibody (anti-rabbit IgG, HRP-linked [7074], Cell Signaling, 1:1000 dilution) for 1 hr at room temperature. The membranes were again washed with TBST, treated with 20X LumiGLO® Reagent and 20X Peroxide (Cell Signaling Technologies #7003) for 1 minute, and imaged for chemiluminescent signal using an ImageQuant Las4010 Digitial Imaging System (GE Healthcare). All signals were normalized to COXIV to account for potential differences in sample loading.

**Preparation of CoA Sepharose Resins**

Synthesis of (Ahx)- and D-(Ahx)-CoA conjugates

The D-(Ahx)_3_-CoA inhibitor scaffold was synthesized on Nova PEG Rink Amide resin via manual peptide synthesis using an orthogonally protected Fmoc-Lys(Boc)-OH. After 3 sequences of 6-(Fmoc-amino)hexanoic acid installation and Fmoc deprotection, an Fmoc-Asp(OtBu)-OH was coupled at the N-terminus, deprotected with 20% piperidine, and coupled with bromoacetic anhydride. The peptide was then cleaved from resin and HPLC purified. The bromoacetamide precursor was then coupled to Coenzyme A (CoA) and purified by preparative HPLC. (Ahx)_4_-CoA inhibitor scaffold was prepared essentially following the same preparation of D-(Ahx)_3_-CoA. Briefly, 4 sequences of of 6-(Fmoc-amino)hexanoic acid installation and Fmoc deprotection were done, followed by a coupling step with bromoacetic anhydride. Peptide was cleaved from resin and HPLC purified. In addition to this manual protocol, we found peptides could also be synthesized by automated peptide synthesis using a PTI Symphony peptide synthesizer, Fmoc- chemistry, HATU coupling reagent, and NovaPEG Ring amide resin. Here lysine residues were introduced using an Fmoc-Lys(Boc)-OH derivative and aspartate residues with Fmoc-Asp(OtBu)-OH. After final Fmoc- deprotection peptide-resin was incubated with bromoacetyl anhydride in DMF (1.5 h), washed with dichloromethane, methanol, and ether, and vacuum dried. Peptides were deprotected/cleaved using 2.5% water, triisopropylsilane in TFA and purified by reverse phase HPLC. In all instances the bromoacetamide precursor was coupled to CoA and purified by preparative HPLC.

Conjugation of (Ahx)- and D-(Ahx)-CoA to Sepharose

(Ahx)_4_-CoA Sepharose and D-(Ahx)_3_-CoA Sepharose were both prepared using NHS-Activated Sepharose 4 Fast Flow Resin essentially following the manufacturer’s protocol (GE Healthcare Life Sciences, Instructions 71-5000-14 AD). Briefly, a 3.4 mM solution of the amine-functionalized capture ligand was prepared in 1x PBS. Then, the Sepharose resin was washed with ice cold 1 mM HCl, pelleted and washing solution removed. The 3.4 mM ligand solution was added at a ratio of 2:1 resin:ligand volume. The pH was then adjusted to ~ 7-8 using 20x PBS before letting the mixture rotate at 4°C overnight. Following the overnight rotation, the resin was pelleted at 1400 rcf for 3 minutes and the supernatant was removed and discarded. Three resin volumes of ice cold 0.1 M Tris-HCl [pH 8.5] was then added to the resin and was left to rotate at room temperature for 3 hrs. The resin was then pelleted again at 1400 rcf for 3 minutes and then washed 3x with ice cold 0.1 M Tris-HCl [pH ~8.5], 3x with ice cold 0.1 M sodium acetate, 0.5 M NaCl [pH 4.5], followed by alternating washes (2x each) between ice cold 0.1 M Tris-HCl [pH ~8.5] and ice cold 0.1 M sodium acetate, 0.5 M NaCl [pH 4.5]. Resin was stored as a 33% solution in aqueous 20% EtOH at 4°C.

**TMT labeling and quantitative LC-MS analysis**

TMT-labeling

Peptides were resuspended in 1mL of 0.1% formic acid (FA) prior to solid phase extraction (Waters SepPak C18, 1mL, 50mg bed volume). Peptides were washed 2x with 0.1% FA prior to elution with 1 mL of 80% acetonitrile (ACN), 0.1% FA. Samples were dried in the speedvac before tandem mass tag (TMT) labeling. Briefly, dried peptides were resuspended in 40 µL 100mM triethylammonium bicarbonate (TEAB) buffer and labeled with 4 channels of the TMT-10plex according to manufacturer protocol (Thermo). The samples were quenched with hydroxylamine to a final concentration of 0.25% for 15 minutes at room temperature. Labeled samples were mixed in equal volumes and dried in the speedvac. Dried samples were stored at -20C until analysis.

Mass spectrometry analysis

Peptides were resuspended in 50 µL buffer A (5% ACN, 0.1% FA) and loaded into autosampler vials. Samples were injected onto the trapping column (Acclaim PepMap C18, 5µm particle, 0.3mm x 5mm) on the Ultimate 3000 liquid chromatography system controlled by Dionex. The peptides were eluted onto an in-house packed separating column with a laser pulled tip (75 µm id, 15cm, 1.9 µm resin (Dr. Maisch)) over 150 minutes with a gradient from 7-40% B (80% ACN, 0.1% FA) before increasing to 60% B in 20 minutes. The column was cleaned by increasing the buffer to 95% for 13 minutes and re-equilibrating for 10 minutes at 2% B to prepare for the next injection. The flow was set to 180 nL/min. The column was directly interfaced with the QExactive Plus mass spectrometer (Thermo) with an applied distal voltage of 2.5 kV. The MS1 spectra were collected at a resolving power of 70,000 with and AGC of 3.0x10^6^. Data dependent MS2 acquisition of the top 15 ions was performed at a resolving power of 35,000 with higher energy collision induced fragmentation at 35% NCE with an isolation window of 0.7 *m/z* and AGC of 1.0x10^5^ as recommended by the instrument vendor.*^1^* Dynamic exclusion was set for 30s.*^2^* Singly charged peptides and those with charges ≥8 were excluded from fragmentation.

Data Analysis

Raw files were analyzed in Proteome Discoverer 2.4 and were searched using SEQEST against a human protein database (downloaded from NCBI 2019-11-03) and a database containing 426 common contaminants with static TMT modification on the peptide N-terminus (+229.163 Da). Variable peptide modifications were added including lysine acetylation (+42.011 Da) or TMT reporter ion modification (+229.163 Da) and methionine oxidation (+15.995 Da). Peptides were normalized on total peptide amount. False discovery rates for the dataset were maintained at 0.006%, 0.01%, 0.023% for peptide spectrum matches, peptides, and proteins, respectively.

Data Accessibility

The mass spectrometry proteomics data have been deposited to the ProteomeXchange Consortium*^3, 4^* via the PRIDE partner repository with the dataset identifier PXD020856 and 10.6019/PXD020856. Reviewer account information:

**Username:**

**Password:** BdHP7ngW

**Chemical capture and immunoaffinity profiling protocol**

Chemoproteomic capture experiments and competitive immunoaffinity profiling were performed essentially as in Montgomery et al.*^5^* Briefly, lysates were first diluted to a working protein concentration of 1.56 mg/mL with ice cold 1x PBS and filtered using a 0.2 µm syringe filter (Celltreat, 229744). Clarified lysate were aliquoted into individual tubes, each tube corresponding to different conditions (i.e. lanes on gel). Lysates were then treated with 20 μL of a cocktail containing either vehicle or competitor ligand (vehicle is 1x PBS for competition experiments where Acetyl-CoA is the competitor ligand, and DMSO for competition experiments where NAA50i [4a] is the competitor ligand) and incubated on ice for 30 minutes. Meanwhile, 33 μL of capture resin (for each condition) was washed three times with 1 mL of ice cold 1x PBS. Competitor/vehicle treated lysates were then transferred to their respective tubes containing capture resin. This mixture was rotated for 1 hr at room temperature, pelleted at 1400 rcf, and supernatant discarded. The Sepharose capture resins were then subjected to a series of mild washes (3 x 500 μl) using ice cold wash buffer (50 mM tris-HCl [pH 7.5], 5% glycerol [omitted in LC-MS/MS experiments], 1.5 mM MgCl_2_, 150 mM NaCl). After the last wash, enriched resin was collected on top of centrifugal filters (MDI, CFPL2101XXXX203). For immunoblot analysis, proteins were eluted from resin via addition of 40 μl 1x SDS sample buffer (Invitrogen, NP0007). Samples were boiled for 10 min at 95°C, and centrifuged at 1400 rcf. Two elutions were performed to maximize KAT isolation. Filtrates from both elutions were combined and loaded onto a 4-12% Bis-Tris gel (Invitrogen, NP0322BOX). After transfer to western blot, membranes were blocked, probed with anti-KAT antibodies [anti-KAT2A (#3305, Cell Signaling Technology, 1:1000 dilution), anti-NAA10 (#13357, Cell Signaling Technology, 1:1000 dilution), anti-ACLY (#4332, Cell Signaling Technology, 1:1000 dilution), anti-NAA25 (ab79490, Abcam, 1:1000 dilution), anti-NAA80 (ab96844, Abcam, 1:1000 dilution), anti-HAT1 (GTX110643, GeneTex, 1:1000 dilution), anti-KAT8 (A300-992A-T, Bethyl Laboratories, 1:1000 dilution), anti-NAA50 (16120-1-AP, ProteinTech 1:1000 dilution)], washed and developed according to the manufacturers protocol. For LC-MS/MS analysis, enriched resins were transferred from centrifugal filters to fresh, 1.7 ml tubes using 400 uL of tryptic digest buffer (50 mM Tris-HCl [Ph 8.0], 1 M Urea). Digests were initiated by the addition of 0.4 μL of 1 M CaCl_2_ and 4 μl of trypsin (0.25 mg/ml), and allowed to proceed overnight at 37°C with shaking. After extraction, tryptic peptide samples were acidified to a final concentration of 55 formic acid and frozen at -80°C for LC-MS/MS analysis.

**Full gels and blots**

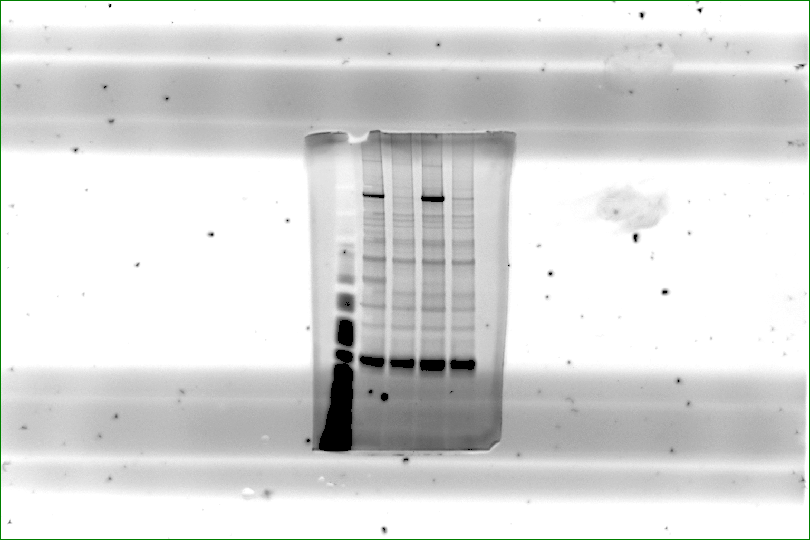

Full blot for Figure 1 and S1. Ladder: MagicMark XP Standard (220 kDa, 120 kDa, 100 kDa, 80 kDa, 60 kDa, 50 kDa, 40 kDa, 30 kDa, 20 kDa).

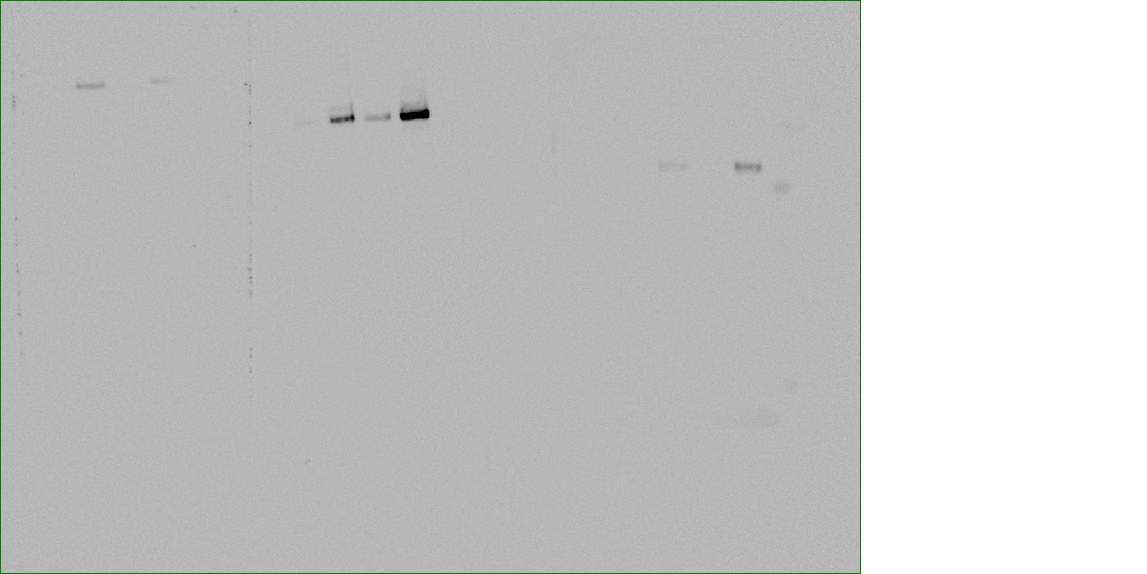

Full blot for Figure 3 and S2. Anti-Gcn5 antibody.

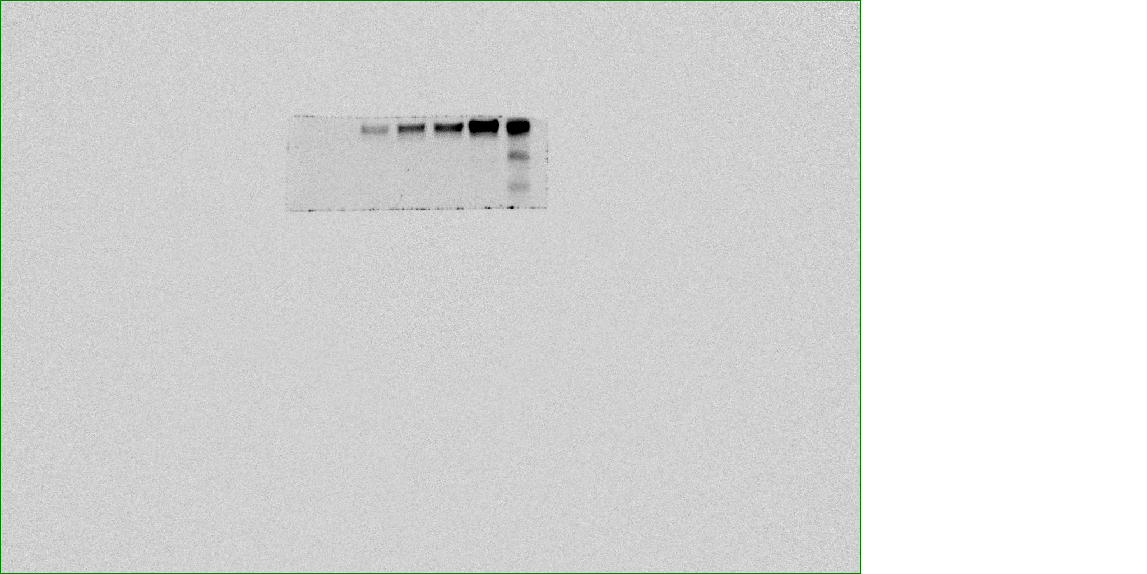

Full blot for Figure 3 and S2. Anti-HAT1 antibody.

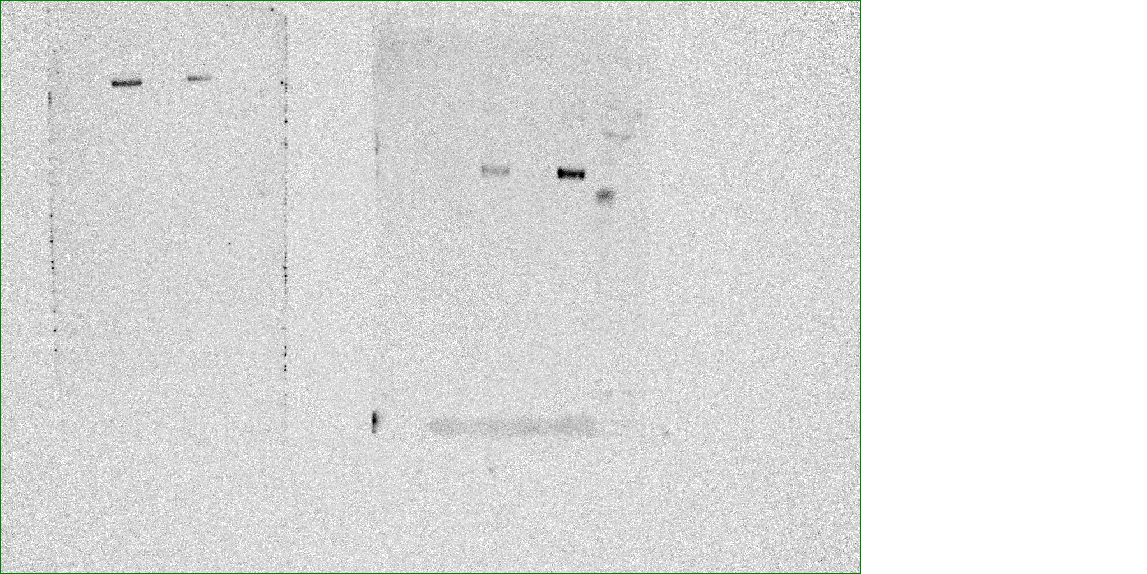

Full blot for Figure 3 and S2. Anti-KAT8 antibody.

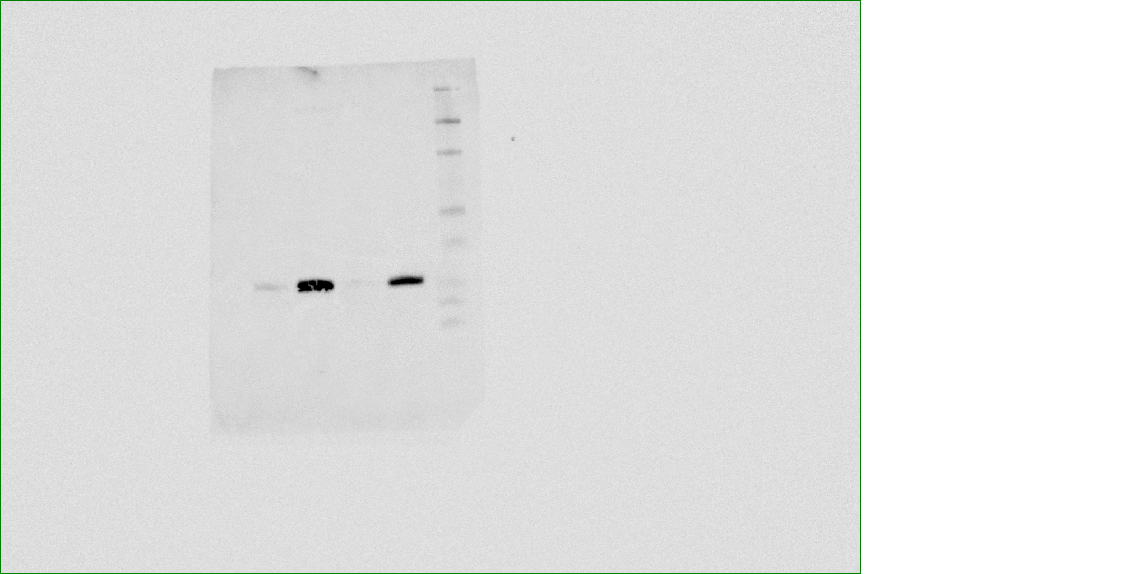

Full blot for Figure 3 and S2. Anti-Naa50 antibody.

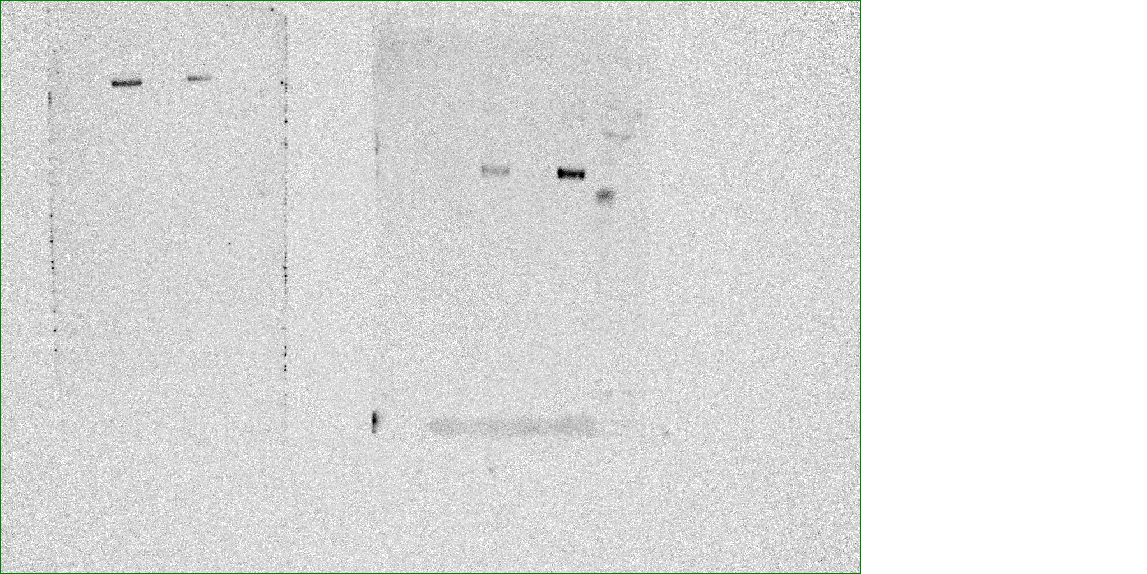

Full blot for Figure 3 and S2. Anti-ACLY antibody.

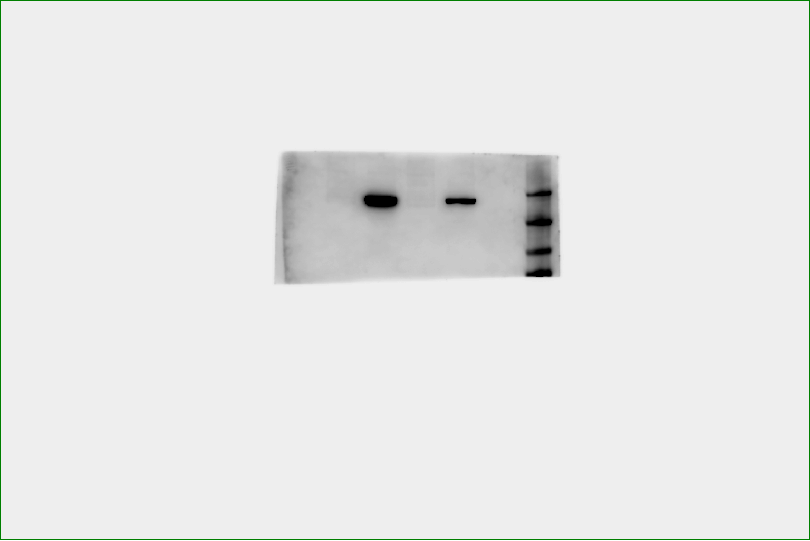

Full blot for Figure 3 and S2. Anti-Naa25 antibody.

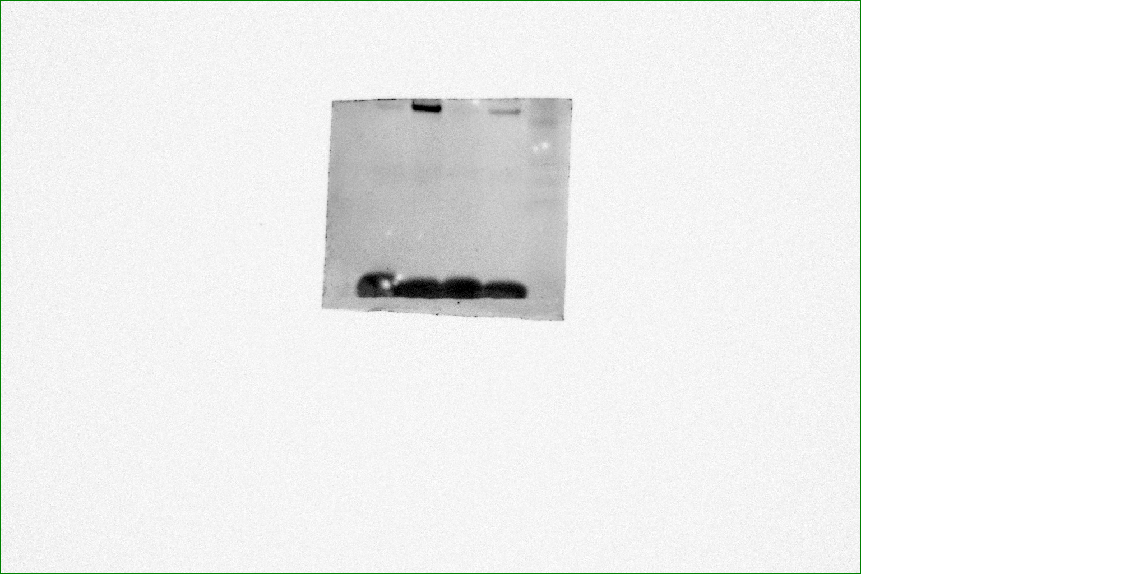

Full blot for Figure 3 and S2. Anti-Naa80 antibody.

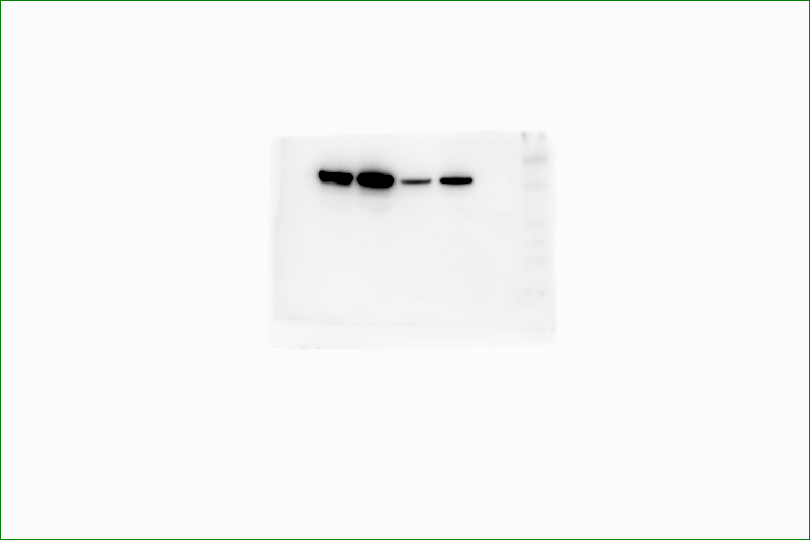

Full blot for Figure 3 and S2. Anti-Naa10 antibody.

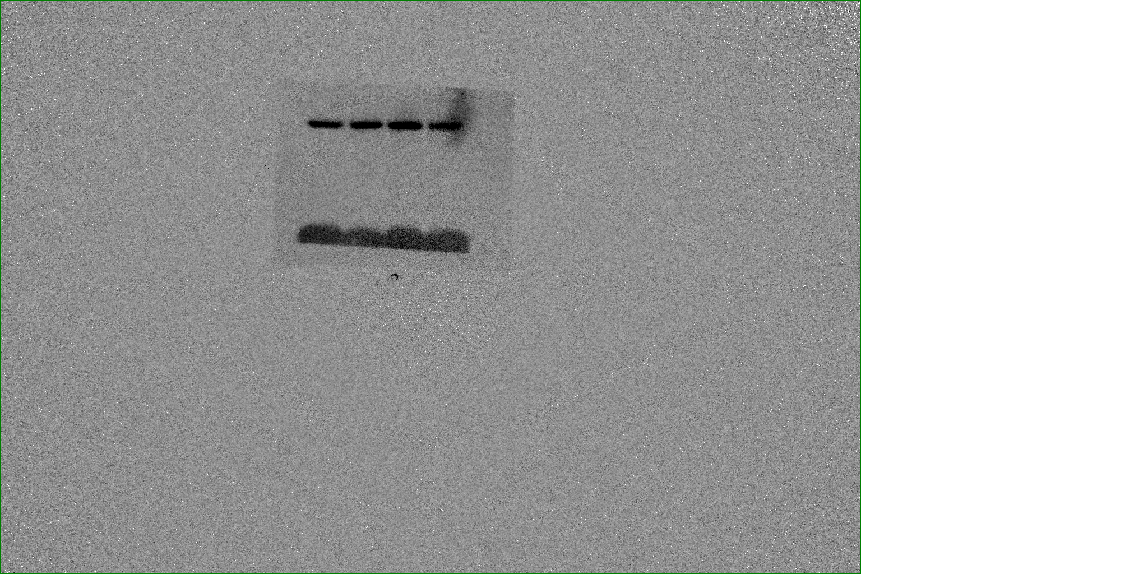

Full blot for Figure 3 and S2. Anti-CoxIV antibody.

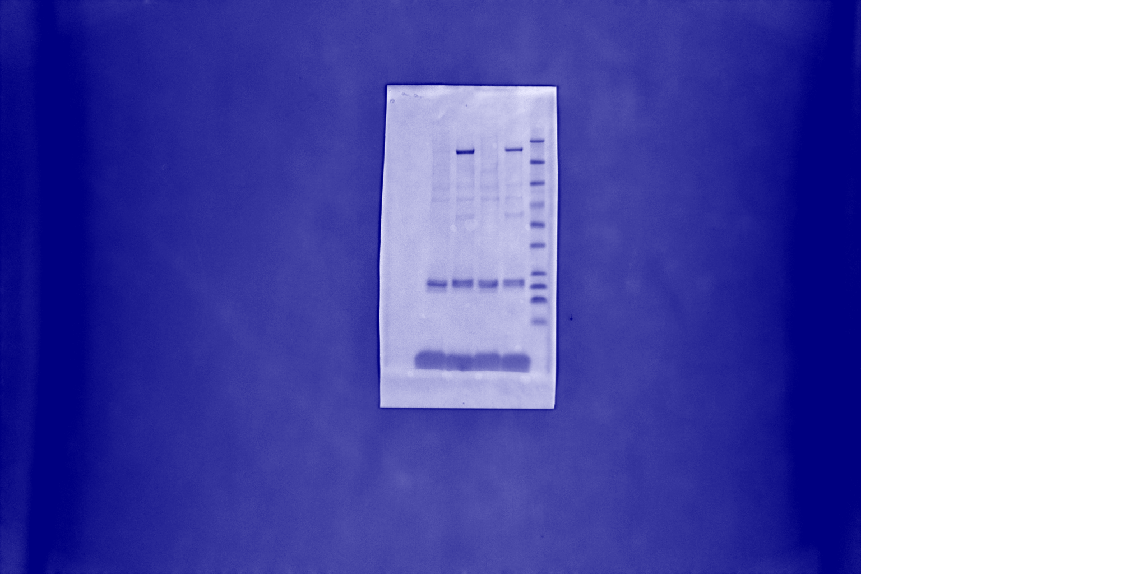

Full blot for Figure 3 and S2. Ponceau stain.

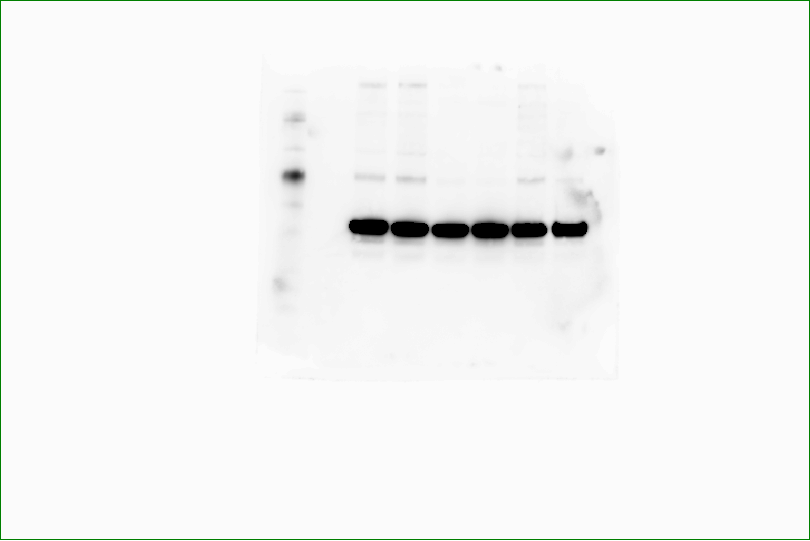

Full blot for Figure 4. Anti-Naa10 antibody.

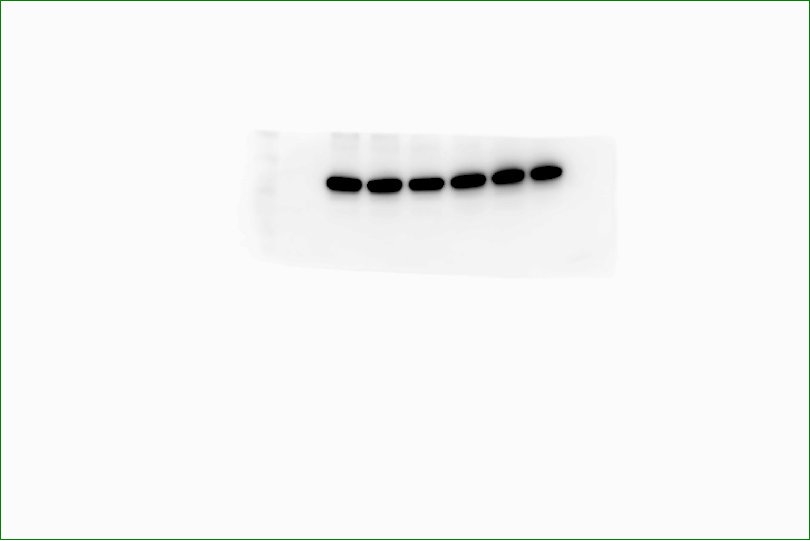

Full blot for Figure 4. Anti-Naa25 antibody.

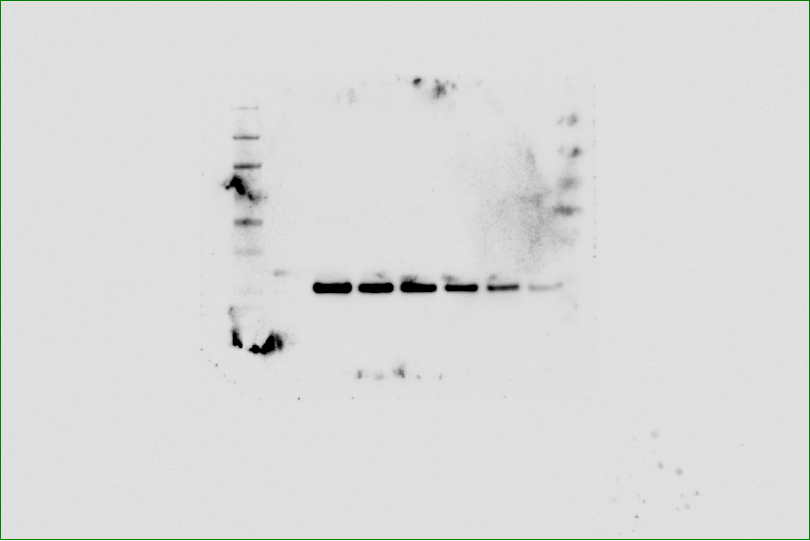

Full blot for Figure 4. Anti-Naa50 antibody.

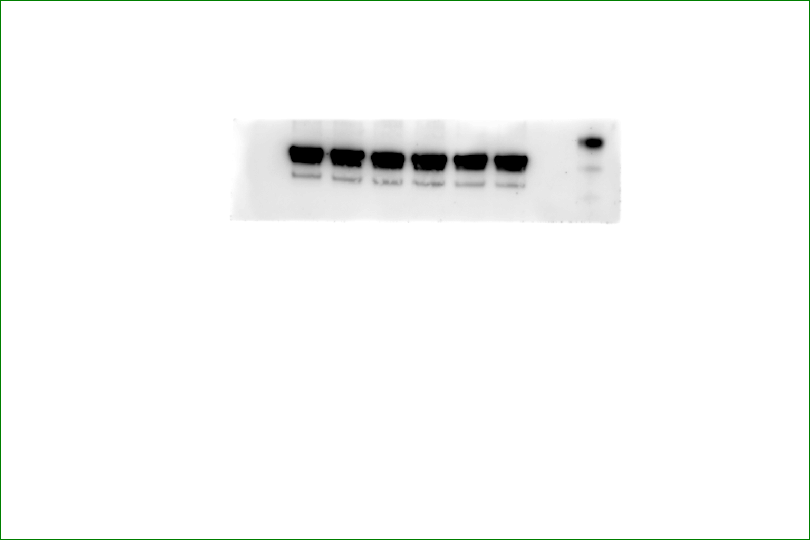

Full blot for Figure 4. Anti-Naa80 antibody.

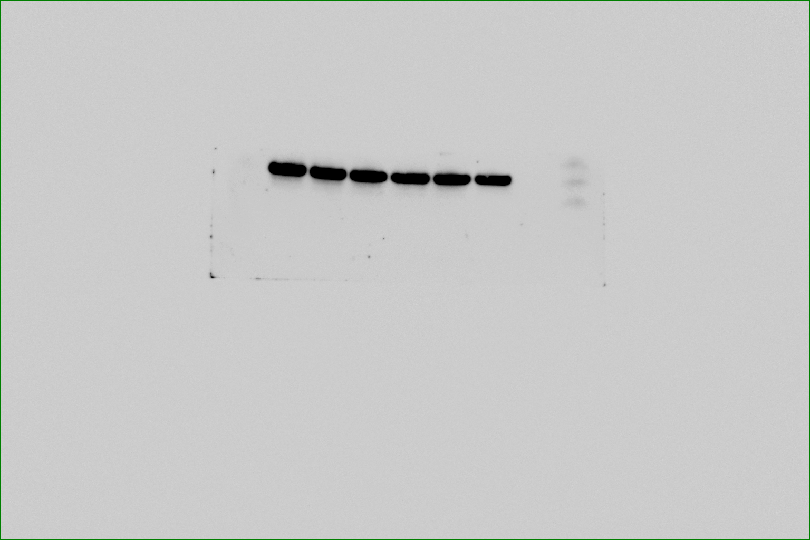

Full blot for Figure 4. Anti-CoxIV antibody.
